# Distributional and Centile Calibration of Diffusion Tensor Imaging Normative Models as Training Sample Sizes Increase to 40,000 Subjects

**DOI:** 10.64898/2026.02.04.703901

**Authors:** Julio E. Villalón-Reina, Yixue Feng, Leila Nabulsi, Talia M. Nir, Sophia I. Thomopoulos, Katherine E. Lawrence, Neda Jahanshad, Seyed Mostafa Kia, Andre F. Marquand, Paul M. Thompson

## Abstract

Normative modeling (NM) is a powerful framework for quantifying individual deviations in brain structure and function relative to a population reference. However, its clinical utility depends on well-calibrated models trained on heterogeneous datasets such as those found in neuroimaging. Here, we systematically examine the effect of training sample size on the distributional and centile calibration of hierarchical Bayesian regression (HBR)-based NMs. Using multisite 3D diffusion MRI scans of the brain from 54,583 subjects, spanning almost the entire lifespan (age: 4-91 years), we trained NMs of white matter fractional anisotropy, a key microstructural metric, on subsamples ranging from 5,000 to 40,000 subjects. HBR was modeled with a Sinh–Arcsinh likelihood. Model calibration was evaluated using Kernelized Stein Discrepancy (KSD) to assess distributional agreement of Z-scores with the standard normal distribution; we also used Mean Absolute Centile Error (MACE) to quantify centile accuracy. Both metrics showed consistent and substantial improvements as the training sample size increased, indicating reduced posterior uncertainty and improved estimation of distributional parameters, particularly at the centile extremes. These results demonstrate that large training cohorts are essential for well-calibrated NMs derived from heterogeneous neuroimaging data and highlight the importance of large-scale data aggregation for reliable individual-level inference.

## I. Introduction

Large-scale international initiatives have greatly expanded access to brain imaging data. This has enabled advanced statistical approaches, such as normative and generative models, to be applied to study imaging-derived brain metrics across different stages of life, health, and disease. Normative modeling (NM) [1], [2] is a statistical framework that estimates the normative distribution - or centiles of variation - of a biological measure within a reference population, based on conditioning variables such as age and sex. This approach allows researchers to detect and monitor brain abnormalities and factors that influence them. Unlike traditional case-control comparisons, NM accounts for individual variability and can even be used to calculate multivariate measures of deviation [3]–[5], offering a more nuanced view of abnormality. As a result, NM has gained considerable attention as it can deepen our understanding of neurological and psychiatric disorders with the goal of supporting personalized medicine.

However, achieving robust and clinically useful normative models requires training on large and diverse populations, as the reference probability distribution must yield accurate centiles when applied to new subjects. This is especially challenging as models often pool heterogeneous data from multiple sites, and also need to model variability in acquisition protocols, demographics, and the relation of normative scores to clinical assessments. Addressing these complexities calls for advanced statistical solutions, such as hierarchical Bayesian regression (HBR), which can model site-specific effects while leveraging shared structure across datasets [6]. These approaches enable scalable and harmonized normative modeling, allowing more accurate detection of individual deviations and improved generalization in clinical settings.

Here we aimed to study the distributional calibration and centile calibration of fitted normative models with an increasing number of participants from 5,000 to 40,000 training subjects (aged 4 to 91 years) using the HBR approach. We expected that larger training sets would reduce posterior uncertainty and would improve estimation of location, scale, and shape parameters of SHASH (the Jones & Pewsey sinh–arcsinh family) distributions [7], [8]. This should theoretically lead to better-calibrated predictive intervals (outer centiles). We demonstrated that increasing the size of the training sample for HBR-NM does indeed lead to better calibration of the empirical distributions and centiles.

Perhaps the closest work to our current report is [9], where the authors evaluated normative modeling methods using simulated hippocampal volume data across sample sizes from 50 to 50,000, focusing on how model choice and sample size affect bias and variance of estimated percentile curves, especially at clinically relevant outer percentiles (e.g., 1st, 5th, 10th). They showed that while variance of percentile estimates decreases with larger samples, residual bias can remain substantial even at large *N*, and uncertainty increased dramatically near the limits of the age range, pointing to challenges in accurately capturing extreme centiles in neuroimaging data. Their results quantify the trade-offs among model flexibility, sample size, and performance, and show that surprisingly large neuroimaging cohorts are needed to reliably estimate clinically important centile curves.

While prior normative modeling work emphasized deviation scores and predictive uncertainty, less attention has been paid to the calibration of estimated centiles, in large, heterogeneous datasets. From a statistical perspective, normative modeling can be viewed as a form of conditional distribution or quantile estimation, connecting it to classical work on quantile regression and modern distributional learning frameworks [10].

## II. Methods

### A. Participants

We included the following public 3D diffusion MRI datasets assessing the brain, with various age ranges to cover the human lifespan: the Adolescent Brain Cognitive Development (ABCD), the Amsterdam Open MRI Collection (AOMIC), the Cambridge Centre for Ageing and Neuroscience (CAMCAN), the Cuban Human Brain Mapping Project (CHBMP), the Chinese Human Connectome Project (CHCP), the Human Connectome Project Aging & Development (HCP_A & HCP_D), the Human Connectome Project Young Adult (HCP_YA), the DTI component of the NIH MRI Study of Normal Brain Development (PedsDTI), the Pediatric Imaging, Neurocognition, and Genetics data repository (PING), the Philadelphia Neurodevelopmental Cohort (PNC), the Queensland Twin Adolescent Brain (QTAB), the Queensland Twin Imaging dataset (QTIM), the Southwest University Longitudinal Imaging Multimodal (SLIM) study, and the UK BioBank (UKBB).

Several clinical datasets were also included: the third phase of the Alzheimer’s Disease Neuroimaging Initiative (ADNI3), the third release of the Open Access Series of Imaging Studies (OASIS3), the Parkinson’s Progression Markers Initiative (PPMI), and the Healthy Brain Network (HBN). Importantly, only the healthy controls (HC) of the latter four datasets were added to the pool of training data to build the normative models, yielding 19 public datasets in total.

From a total of 54,583 individuals (age: 4-91 years), we subsampled six times to yield training samples of 5,000, 10,000, 15,000, 20,000, 30,000 and 40,000 subjects, with random subsampling 10 times with different seeds.

All subjects were processed according to the ENIGMA-DTI protocol [11]. As an illustrative sample, the average of the fractional anisotropy across the entire skeleton of the ENIGMA-DTI template was extracted for all participants.

For illustrative purposes we show a plot of the average WM normative model across the lifespan [6] (**Figure 1**).

**Fig. 1.**
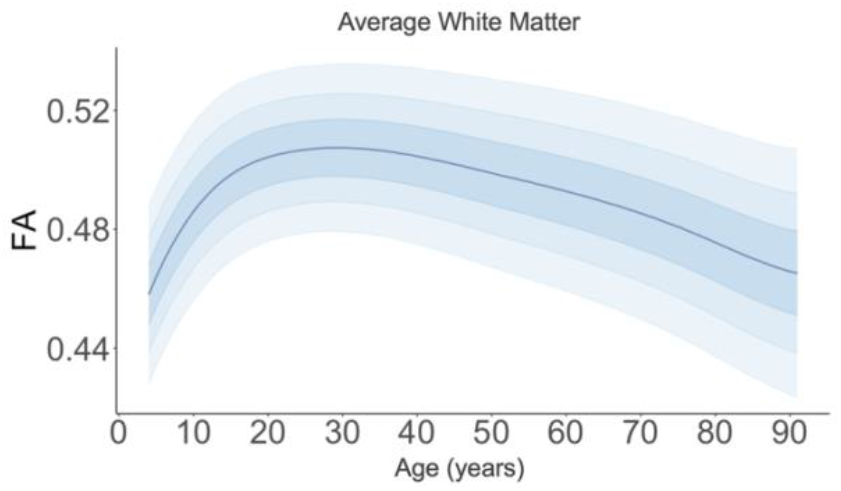
Normative model of the average fractional anisotropy (FA) of the entire white matter region of the brain across the lifespan from 4 to 91 years. The plot shows the median and the centile curves for Z=±0.68, ±1.29 and ±2.

All study participants had information on age and sex at the time of the imaging session, along with a usable dMRI scan. ADNI3, OASIS3, PPMI, CAMCAN and HBN datasets were manually checked for head movement, FOV (field of view) related artifacts and other dMRI acquisition artifacts and preprocessing errors caused by poorly tuned software tools. For all other datasets used for training, outliers were detected with an anomaly detection algorithm called isolation forests implemented in *Scikit-learn*.

We then ran HBR with the ‘average’ FA as the response variable for each subject, age as the explanatory variable and we modeled as a batch effect the site (n=37 values) and sex (2 possible values). We included a random intercept for each batch effect: site and sex.

We modeled the output response, i.e., the ROI per DTI metric, using a SHASH likelihood [7]. The standard SHASH distribution arises by applying a sinh–arcsinh transformation to a normal distribution (sinh-arcsinh Distribution) yielding a 4-parameter family (*µ*,σ,ε,δ) parameterized by location, scale, skewness (ε), and tail weight (δ). If *Z* ∼*N*(0,1), we can define the sinh–arcsinh transformation where *µ* ∈ℝis the location parameter, σ>0is the scale, ε ∈ℝcontrols skewness, and δ> 0 controls tail weight, as follows:

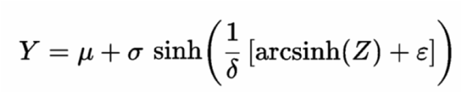

The resulting random variable *Y* follows the SHASH distribution, yielding a flexible four-parameter family capable of modeling asymmetric and heavy-tailed deviations from Gaussianity.

We ran HBR for each training sample (6 total), 10 times for each sample.

### B. HBR evaluation

To evaluate the calibration of our hierarchical Bayesian regression (HBR) models, we computed the kernelized Stein discrepancy (KSD) between the empirical Z-scores obtained from HBR and a standard normal distribution [12].

The KSD is based on Stein’s identity, which compares an empirical distribution *q* to a target distribution *p* using the *score function:*

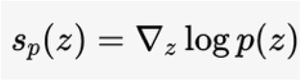

which for the standard normal simplifies to *s*_*p*_ (*z*) = −*z*. This formulation allows discrepancies to be detected without requiring normalization of the target density. KSD is especially well-suited to understanding the distributional agreement between a sample of points and a reference model, and it can be computed if one of the data densities is only available in the form of a sample. If *z*_*i*_, for *i* = 1 to *N*, denotes the sample of standardized Z-scores, KSD is a nonparametric measure of distributional discrepancy: smaller values indicate closer agreement with the expected 𝒩(0,1) distribution, whereas larger values reflect deviations such as skewness or heavy tails.

KSD is a practical variant of the Stein discrepancy that leverages reproducing kernels to make computation tractable. Using a positive-definite reproducing kernel *k*(⋅,⋅), typically an RBF kernel, the squared KSD between the empirical distribution of Z-scores and the target normal distribution can be estimated as:

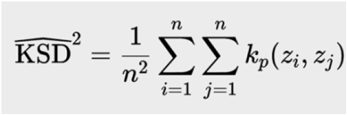

- where *k*_*p*_ (⋅,⋅) is the kernel induced by applying the Langevin–Stein operator to *k*. A near-zero KSD implies that the HBR-derived Z-scores align well with the theoretical *k*expectation, while higher values may indicate model misspecification or residual heterogeneity. This formulation allows KSD to compare the entire empirical distribution to the target without requiring normalization of the target density. By doing so, KSD provides a rigorous way to assess whether the fitted model produces Z-scores consistent with the theoretical expectation under Normal(0,1). In our analysis, we report the square root of the KSD statistic, where lower values reflect better calibration.

We also used the mean absolute centile error (MACE), a metric proposed by Zamanzadeh et al. [13], to evaluate the centiles. This is a metric designed to directly assess centile calibration in normative models. It measures how well the predicted centiles (e.g., 1st, 5th, 25th, 50th, 75th, 95th, 99th) align with the empirical centiles observed in the test data. Intuitively, if a model predicts that a subject falls at the 5th centile, then across many test subjects, close to 5% should actually fall below that predicted value. MACE computes the average absolute difference between the predicted centile and the observed proportion of data below that threshold. A lower MACE implies a better calibration, where predicted centiles match observed data well; higher values indicate miscalibration (e.g., systematic bias, poor uncertainty modeling).

## III. Results

**Figure 2A** shows the gradual decline in KSD values as we increase the sample size used to train the HBR normative model. Each data point represents each of the 10 fitted normative models (10 different iterations). For each training sample, we show the 2.5 to 97.5-th percentiles. The line connects the median KSD value for each training sample size.

**Fig. 2.**
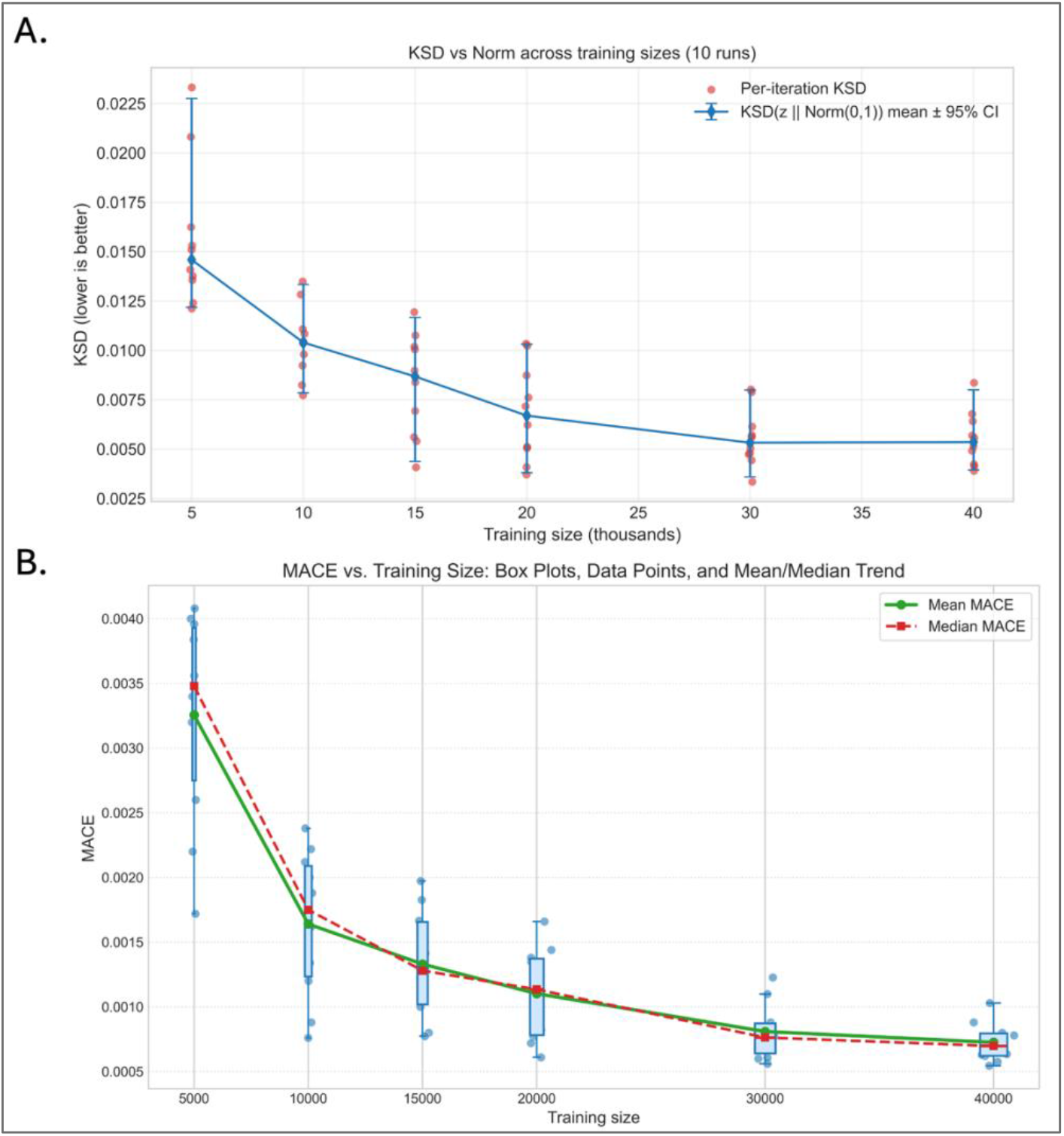
Effect of training sample size on calibration of hierarchical Bayesian normative models. **(A) Kernelized Stein Discrepancy** (KSD) between HBR-derived Z-scores and the standard normal distribution, shown as a function of training sample size (5k–40k). *Red points* indicate per-iteration KSD values across 10 random subsampling runs at each sample size. *Blue markers* and the connecting line show the mean KSD, with error bars denoting the 95% confidence interval across runs. Lower KSD values indicate better distributional calibration, with a clear monotonic improvement as training size increases. **(B) Mean Absolute Centile Error** (MACE) as a function of training sample size, summarizing centile calibration across the same 10 runs. *Box plots* show the distribution of MACE values at each sample size, and overlaid points represent individual runs. The *green solid line* denotes the mean MACE, and *the red dashed line* denotes the median MACE. Both mean and median MACE decrease steadily with increasing training size, indicating improved alignment between predicted and empirical centiles at larger cohort sizes.

**Figure 2B** shows the MACE values when increasing the training sample used for HBR normative modeling. We show the box plot for the MACE values for each training sample size.

Figure 3. shows the centiles (0.05, 0.25, 0.5, 0.75, 0.95) of the fitted normative models with increasing sample sizes, for just one random subsampling iteration (out of 10). The blue data points correspond to the females of the ABCD dataset. All “other data” points in black correspond to all sites that are not ABCD. Age is in years is divided by 100.

**Fig. 3.**
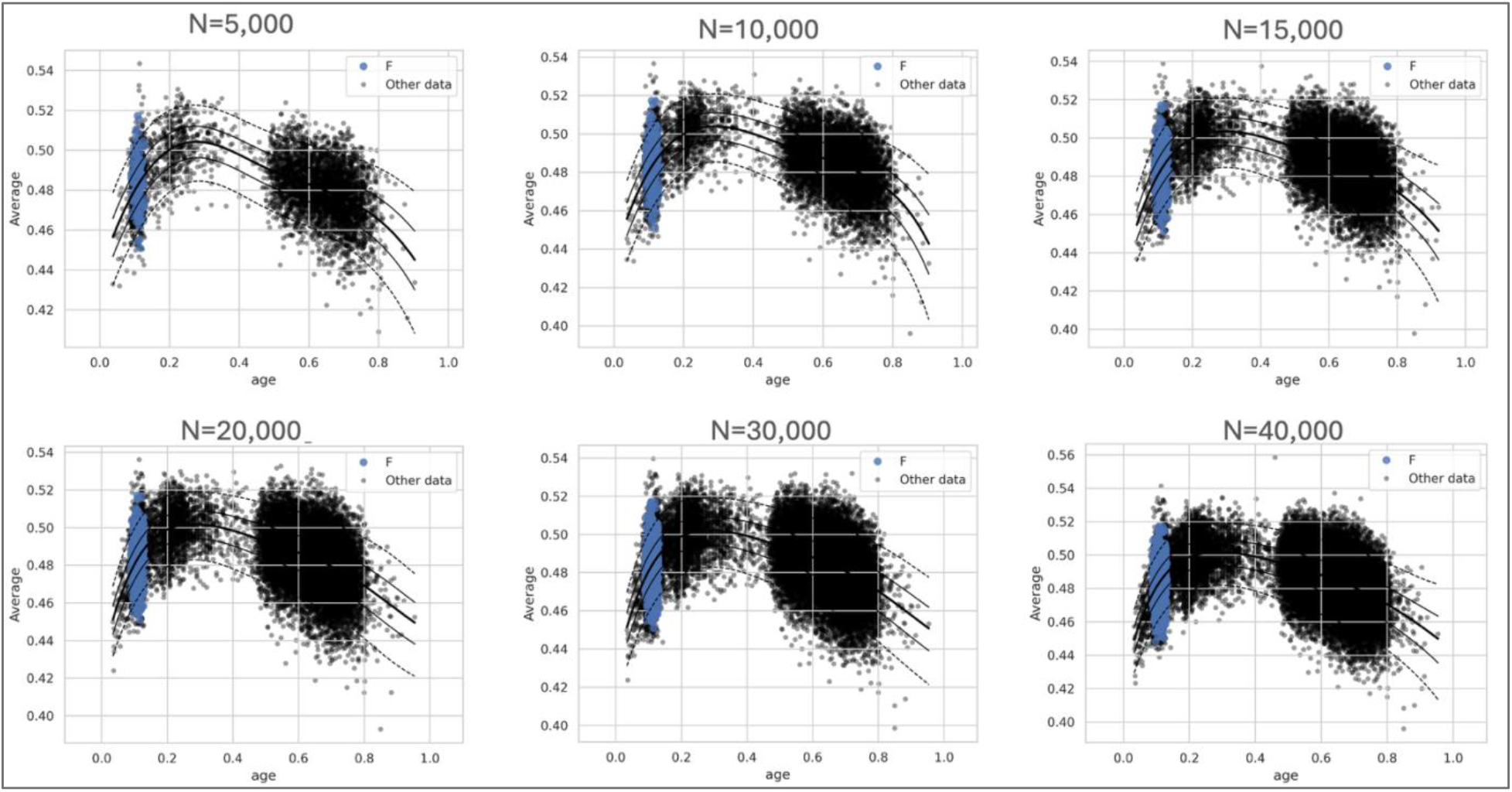
Effect of training sample size on estimated normative centiles across the lifespan. Normative trajectories of average white matter fractional anisotropy (FA) as a function of age are shown for hierarchical Bayesian regression models trained on increasing sample sizes (N = 5,000 to 40,000). Black points represent data from all sites except ABCD (“Other data”), while blue points correspond to female participants from the ABCD cohort, shown for reference. Solid black curves denote fitted centiles (5th, 25th, 50th, 75th, and 95th), with dashed curves indicating the outer centiles. As the raining sample size increases, centile estimates become smoother and more stable across age, with reduced curvature variability and tighter separation between adjacent quantiles. Larger samples yield more consistent estimation of the tails, reflecting improved identifiability of higher-order distributional parameters (skewness and tail weight) under the sinh–arcsinh likelihood. These trends show how increased sample size enhances the numerical stability of quantile estimation and the effective uniform convergence of empirical to model-based centiles across the age range.

The reduction of calibration error with sample size motivates the use of large samples to build accurate normative models. The observed monotonic improvement in centile calibration is consistent with classical uniform convergence results for empirical distribution functions, suggesting that theoretical sample-size guarantees for normative centiles may be attainable. Future work will investigate theoretical laws that govern sample efficiency of these models, including the Dvoretzky–Kiefer– Wolfowitz inequality and its extension to multivariate distributions [14].

## IV. Conclusions

We found that increasing training sample size markedly improves both distributional and centile calibration of hierarchical Bayesian normative models for diffusion MRI, with the largest gains observed at the distributional extremes that are most relevant for clinical inference. Using kernelized Stein discrepancy and mean absolute centile error, models trained on up to 40,000 subjects yielded more stable estimates of location, scale, skewness, and tail-weight parameters under a flexible sinh-arcsinh likelihood, reducing posterior uncertainty and centile miscalibration. Improvements with increasing sample size are most pronounced at the outer centiles, consistent with the fact that tail parameters in flexible distributions (e.g., SHASH) are weakly identifiable in small samples. The median trajectory (50^th^ percentile) stabilizes at relatively small sample sizes, even when centile calibration continues to improve substantially, indicating that accurate mean modeling alone is insufficient for reliable individual-level inference.

These empirical findings motivate several important extensions. The reduction in centile error with sample size suggests that finite-sample convergence guarantees for normative centiles may be derived using tools from empirical process theory, such as the Dvoretzky-Kiefer-Wolfowitz inequality and related uniform convergence results. Establishing explicit sample-efficiency bounds for calibrated centile estimation would provide principled guidance on the cohort sizes needed for reliable individual-level inference, particularly for extreme quantiles.

An additional direction for future work could be the explicit modeling of scanning protocol effects. While hierarchical Bayesian regression accounts for site-level shifts, acquisition parameters may also influence variance, skewness, and tail behavior of diffusion measures. Incorporating scanner and protocol characteristics as structured covariates - affecting the mean and higher-order distributional parameters - may further improve calibration and reduce residual heterogeneity in multi-site normative models. It is likely that acquisition protocol effects may act on variance, skewness, or tail weight rather than on the mean alone.

From a statistical perspective, normative modeling can be viewed as conditional distribution estimation rather than mean regression. While prior work largely focused on deviation scores, our results emphasize the importance of centile calibration itself, particularly at the distributional extremes. The marked reduction in tail instability with increasing sample size suggests improved identifiability of higher-order distributional parameters, such as skewness and tail weight, which are weakly constrained in smaller cohorts. These findings align with classical results on uniform convergence of empirical distributions and motivate future work on theoretical sample-efficiency guarantees for normative centiles. Although demonstrated here for a univariate white matter diffusion metric, these results are likely to generalize to multivariate and high-dimensional normative modeling frameworks. As normative models scale to richer representations of brain structure and function, rigorous calibration assessment will be essential for ensuring robustness and clinical utility. Flexible distributional modeling and principled calibration metrics will be crucial for advancing the practicality of normative modeling for precision neuroscience.

